# Exosome-mediated apoptosis pathway during WSSV infection in crustacean mud crab

**DOI:** 10.1101/2020.02.03.932970

**Authors:** Yi Gong, Tongtong Kong, Xin Ren, Jiao Chen, Shanmeng Lin, Yueling Zhang, Shengkang Li

## Abstract

MicroRNAs are regulatory molecules that can be packaged into exosomes to modulate recipient’s cellular response, while their role during viral infection is beginning to be appreciated. However, the involvement of exosomal miRNAs during immunoregulation in invertebrates has not been addressed. Here, we found that exosomes released from WSSV-injected mud crabs could suppress viral invasion by inducing apoptosis of hemocytes. Besides, miR-137 and miR-7847 were found to be less packaged in mud crab exosomes during viral infection, with both miR-137 and miR-7847 shown to be negative apoptosis regulators by targeting the apoptosis-inducing factor (AIF). Moreover, our data revealed that AIF did not only translocate to the nucleus to induce DNA fragmentation, but could also competitively bind to HSP70 to disintegrate the HSP70-Bax (Bcl-2-associated X protein) complex, which eventually activated the mitochondria apoptosis pathway via free Bax. Therefore, our findings provides a novel mechanism underlying the crosstalk between exosomal miRNAs and apoptosis pathway in innate immunity in invertebrates.

## Introduction

Exosomes, measuring 30-120 nm in diameter, are extracellular vesicles of endocytic origin that are released into the extracellular environment under physiological and pathological conditions [1, 2]. Exosomes can be produced by various types of donor cells and then transferred to target cells, which serve as mediators during intercellular communications by transporting information cargo, including lipids, proteins, mRNAs and microRNAs (miRNAs) [3, 4]. Specific proteins highly enriched in exosomes are usually used as markers to identify exosomes, such as TSG101, CD9, CD63 and CD81 [5, 6]. As a form of intercellular vesicular transport, exosomes are involved in the regulation of a variety of pathological processes [7]. Recently, exosomes have been implicated in viral pathogenesis and immune responses [8, 9]. During virus infection, the infected host cells can excrete exosomes containing viral or host genetic elements to neighboring cells to help modulate host immune response [10, 11], which suggest a crucial role of exosomes during virus infection. However, very little is known about how exosomes regulate host immune response and impact on viral infection.

miRNAs are small non-coding RNAs of 18-25 nucleotides in length that can bind to the 3’-untranslated region (UTR) of target genes in most cell types [12, 13]. Binding of the miRNAs can lead to recruitment of the target mRNAs to RNA-induced silencing complex (RISC), which result in translational arrest or mRNA degradation and decreased protein expression of the target genes [14, 15]. Apart from their endogenous actions, miRNAs can be secreted into the extracellular space within exosomes, with these miRNA-containing exosomes being taken up into neighboring or distant cells to modulate the expression of multiple target genes in the recipient cells [16, 17]. RNA sequencing analysis has shown that miRNAs are the most abundant among the exosomal RNA species [18]. Recent evidences indicate that the alteration of miRNA composition can significantly affect the biological activities of exosomes that have been taken-up during virus infection [19, 20]. Importantly, it has been demonstrated that package of miRNAs into exosomes is selective and can also reflect the dysregulated miRNA composition in donor cells [21]. It is conceivable that exosome-mediated intercellular transfer of miRNAs contributes to immune defense of the recipient cells and regulate viral spread.

White spot syndrome virus (WSSV) is a large enveloped double-stranded DNA virus that causes huge economic losses in aquaculture [22]. In recent years, studies have shown the widespread pathogenicity of WSSV among many marine crustaceans, including shrimp, crayfish and crabs [23]. Generally, innate immune responses (humoral and cellular), are used by invertebrates to recognize and protect themselves against pathogenic microbes [24]. Apoptosis is one type of cellular immune response that plays an essential role in host antiviral immunity [25], with viral infection capable of inducing apoptosis in infected cells in both vertebrates and invertebrates [26]. Since exosomes are widely thought to be effective defense tools for host resistance to viral infection [27], while marine crustaceans possess an open circulatory system, makes this an ideal carrier for exosomes to perform their immune functions. However, the role of exosomes during antiviral immune response in marine crustaceans is uncler, while whether there is the involvement of apoptosis remains unknown.

In an attempt to explore the involvement of exosomes in apoptosis during antiviral immunoregulation of marine crustaceans, the interactions between exosomal miRNAs and WSSV infection were characterized in mud crab *Scally paramamosain*. The results revealed that exosomes released from WSSV-injected mud crabs could suppress viral invasion by inducing hemocytes apoptosis. Moreover, it was found that miR-137 and miR-7847 were less packaged in exosomes after WSSV challenge, resulting in the activation of AIF, which eventually caused apoptosis and suppressed viral infection of the recipient hemocytes.

## Results

### The involvement of exosomes in antiviral regulation of mud crab

To characterize exosomes from mud crab during WSSV infection, exosomes (i.e., exosome-PBS and exosome-WSSV), isolated from hemolymph of healthy (PBS-injected) and WSSV-injected mud crabs, respectively were used. The cup-shaped structure and size of the isolated exosomes were detected by electron microscopy (Fig. 1A) and Nanosight particle tracking analysis (Fig. 1B). In addition, Western blot analysis of exosome markers (CD9 and TSG101) and cytoplasmic marker (calnexin) were further used to ascertain that the isolated particles were exosomes (Fig. 1C). Furthermore, to analyze the capacity of the isolated exosomes to be internalized by hemocytes, mud crabs were injected with exosomes labeled with DiO (green), and hemocytes then isolated and labeled with Dil (red). Confocal microscopic observation showed that the isolated exosomes (from PBS and WSSV injected mud crabs) were successfully internalized in hemocytes (Fig. 1D). Besides, the effects of the obtained exosomes on WSSV proliferation was investigated using real-time PCR analysis. It was observed that mud crabs injected with exosome-PBS had a significantly higher WSSV copy number than those injected with exosome-WSSV (Fig. 1E). These results suggest that the secreted exosomes could be internalized in hemocytes, which might play an important role in the immune response of hemocytes in mud crab against virus infection.

**Fig 1.**
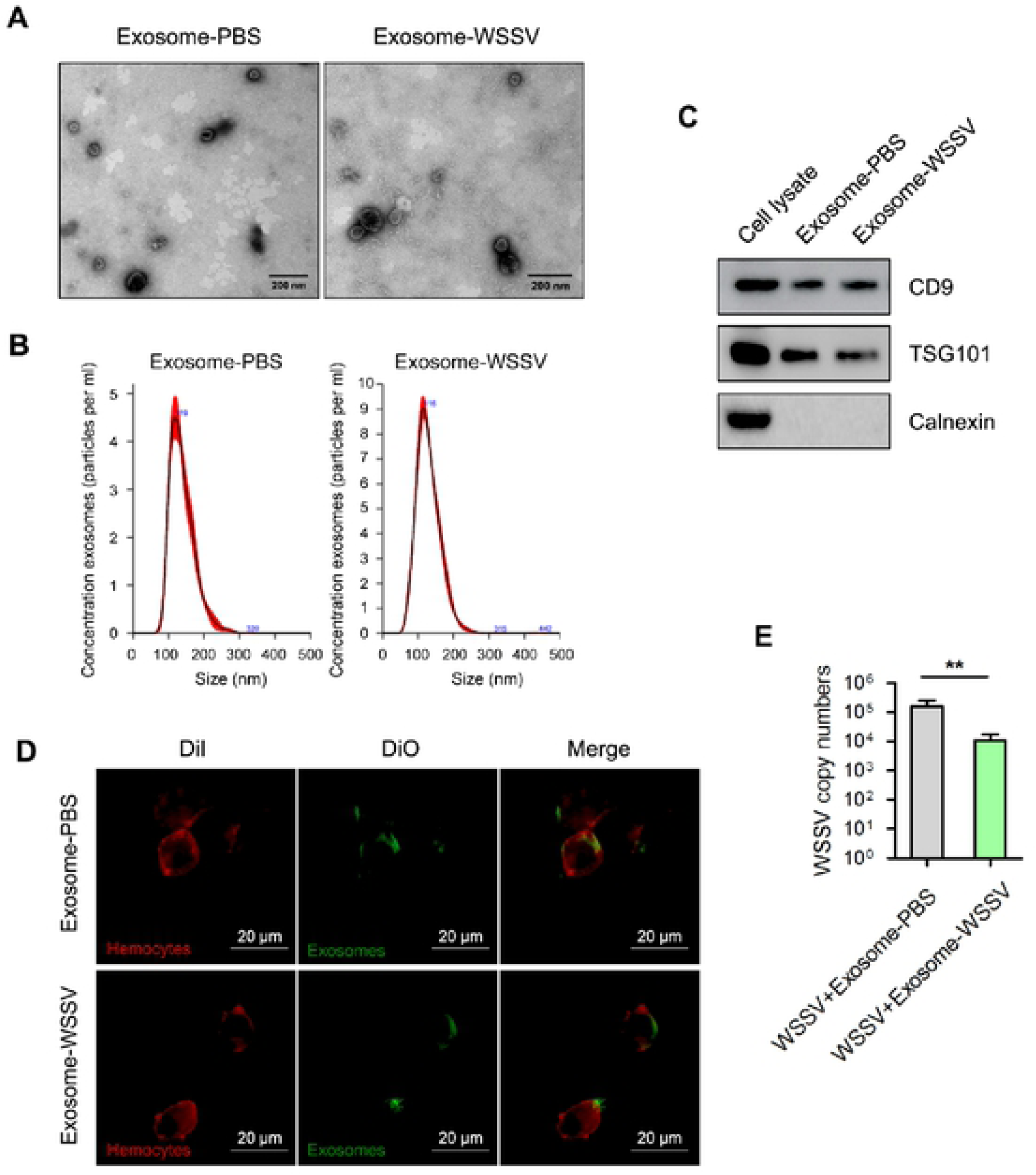
Exosomes secreted from WSSV-infected mud crab participate in antiviral regulation. **(A-B)** Exosomes isolated from mud crab with different treatments were detected by electron microscopy **(A)** and Nanosight particle tracking analysis **(B)**. Scale bar, 200 nm. **(C)** Western blotting assay of exosomal marker proteins (CD9 and TSG101) and cytoplasmic marker protein Calnexin in cell lysate and exosomes. **(D)** Confocal imaging showed the delivery of Dio-labeled exosomes (green) to Dil-labeled mud crab hemocytes (red). Scale bar, 20 μm. The indicated exosomes were injected into mud crab for 6 h, then the hemocytes were isolated and subjected to confocal imaging analysis. **(E)** The involvement of exosomes during WSSV infection, mud crabs were co-injected with the indicated exosomes and WSSV for 48 h, followed by the detection of WSSV copies. All data represented were the mean ± s.d. of three independent experiments (**, *p*<0.01).

### Exosome-mediated viral suppression is relevant to apoptosis activation

To investigate the involvement of apoptosis during exosome-mediated virus suppression in mud crab, exosome-PBS or exosome-WSSV and WSSV were co-injected into mud crabs. The number of apoptotic hemocytes (at 48 hpi) were evaluated by Annexin V/PI staining. The results showed higher apoptosis in the exosome-WSSV and WSSV co-injected group compared with the other groups (control and exosome-PBS groups) (Fig. 2A). Caspase 3/7 activity was found to significantly increase in the exosome-WSSV and WSSV co-injected group compared to the control group (Fig. 2B). To better understand the role of exosomes in mediating mitochondrial membrane potential, mud crabs were co-injected with either exosome-PBS or exosome-WSSV and WSSV, and hemocytes analyzed using confocal microscopy. The confocal microscopic observation revealed that the mitochondrial membrane potential showed weak red fluorescence (based on JC-1 aggregates) and strong green fluorescence (JC-1 monomers) in both exosome-injected groups, compared with the controls (Fig. 2C). Moreover, the proapoptotic protein, BAX, was upregulated, while prosurvival Bcl-2 was downregulated in hemocytes of exosome co-injected mud crabs compared with controls (Fig. 2D). To reveal the interactions between apoptosis and virus infection in mud crab, the apoptosis inducer cycloheximide and apoptosis inhibitor Z-VAD-FMK were used to evaluate their effects on WSSV proliferation in hemocytes of mud crab. The results showed significantly lower WSSV copy number in the cycloheximide and WSSV-injected group, but significantly higher WSSV copy number in the Z-VAD-FMK and WSSV-injected group, compared with the WSSV-injected group (Fig. 2E), which suggest a negative role of apoptosis during virus infection. To further investigate the effect of apoptosis on exosome-mediated virus suppression, mud crabs were co-injected with WSSV and either exosome-PBS, exosome-WSSV, or exosome-WSSV and Z-VAD-FMK. It was found that the exosome-WSSV-mediated virus suppression was significantly weaken when apoptosis was inhibited by Z-VAD-FMK (Fig. 2F). These results revealed that exosomes isolated from WSSV challenged mud crabs suppressed viral infection through activation of apoptosis.

**Fig 2.**
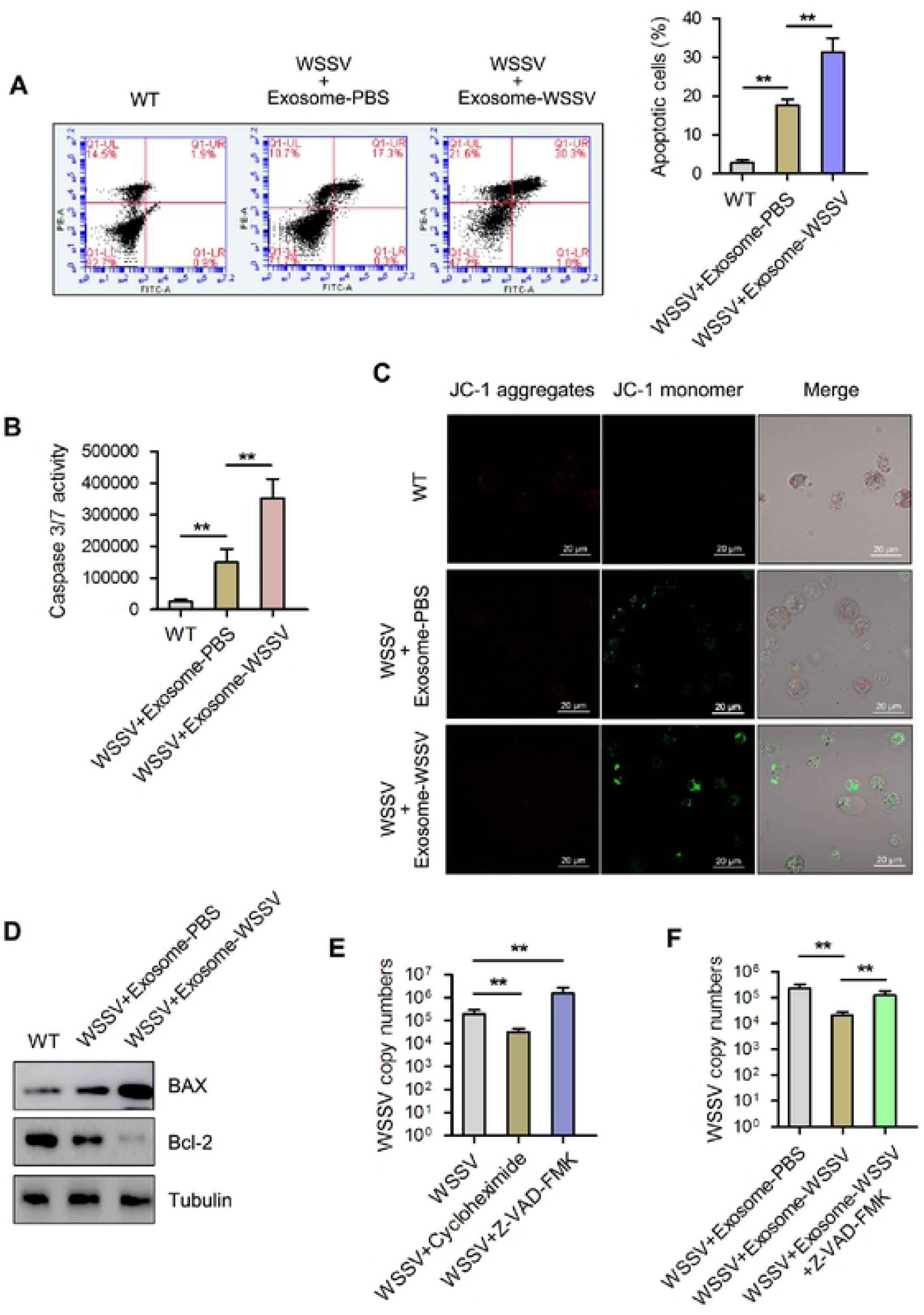
Exosomes isolated from mud crab challenged with WSSV suppressed virus infection through activation of apoptosis. **(A-D)** The influence of the indicated exosomes on apoptosis of mud crab hemocytes. The isolated exosomes from mud crab with different treatments (including PBS and WSSV) were co-injected with WSSV into mud crab for 48 h, then, the apoptotic levels of the hemocytes were examined through annexin V assay **(A)**, caspase 3/7 activity analysis **(B)**, mitochondrial membrane potential measurement **(C)** and apoptosis-associated protein detection **(D)**. **(E)** The role of apoptosis regulation during virus invasion, apoptosis inducer cycloheximide or apoptosis inhibitor Z-VAD-FMK were co-injected into mud crab with WSSV for 48 h, followed by the detection of WSSV copies. **(F)** The involvement of apoptosis regulation during exosome-mediated virus suppression, mud crabs were co-injected with the indicated exosomes, WSSV and apoptosis inhibitor Z-VAD-FMK, then WSSV copy numbers were detected. All the data were the average from at least three independent experiments, mean ± s.d. (**, *p*<0.01).

### Functional miRNA screening in exosomes

Microarray analysis of exosomal miRNAs was carried out with a 1.5-fold change and *P<0.01* used as threshold cut-off. The results showed that 124 miRNAs were found to be differentially expressed in isolated exosomes released from the exosome-WSSV group compared with the exosome-PBS group. Among the differentially expressed miRNAs, 84 were upregulated and 40 were downregulated (see heatmap in Fig. 3A). The top 10 differentially expressed miRNAs, including miR-137, miR-60, miR-373, miR-7847, miR-87a, miR-513, miR-353, miR-81, miR-508, and miR-387, are shown in Fig. 3A. To investigate whether these miRNAs affect WSSV proliferation, miRNA mimics (miR-137 mimic or miR-7847 mimic) or anti-miRNA oligonucleotides (AMOs) (AMO-miR-137 or AMO-miR-7847) were co-injected with WSSV into mud crabs for 48 hours. The results revealed an increase in WSSV copy number in mud crabs after miRNA mimics injection and a decrease in WSSV copy number in mud crabs after AMO injection (Fig. 3B and 3C). The expression levels of miR-137 and miR-7847, normalized to U6, were investigated in exosome-PBS and exosome-WSSV injected mud crabs. It was found that both miR-137 and miR-7847 were significantly downregulated in the exosome-WSSV injected group, compared to the exosome-PBS injected group (Fig. 3D).

**Fig 3.**
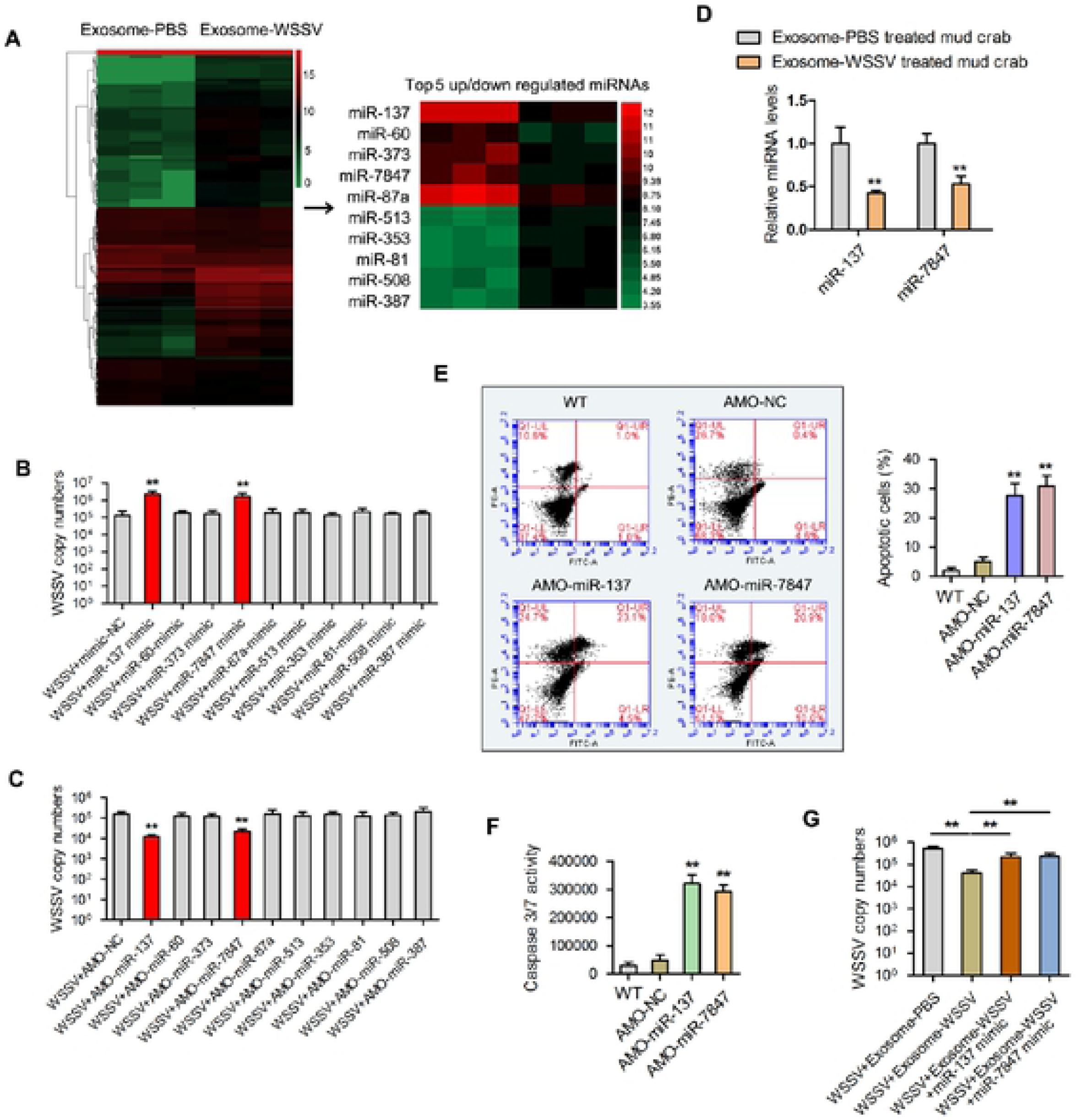
Exosomal miR-137 and miR-7847 were characteristically secreted to mediate apoptosis and virus invasion in mud crab. **(A)** Microarray analysis of exosomal miRNAs were presented in a heatmap, the top5 up/down regulated miRNAs in the indicated exosomes were listed in detail. **(B-C)** The effects of the indicated miRNAs on virus infection, mimics or anti-miRNA oligonucleotides (AMOs) of the indicated miRNAs were co-injected with WSSV into mud crab for 48 h, then WSSV copy numbers were evaluated via qPCR. **(D)** The expression levels of miR-137 and miR-7847 in mud crab challenged with different exosomes. **(E-F)** The functions of miR-137 and miR-7847 on apoptosis regulation, AMO-miR-137 and AMO-miR-7847 were injected into mud crab separately, then the hemocytes were subjected to annexin V assay **(E)** and caspase 3/7 activity analysis **(F)**. **(G)** The participation of miR-137 and miR-7847 during exosome-mediated virus suppression, the indicated exosomes, WSSV and AMOs were co-injected into mud crabs, followed by the detection of WSSV copies. Experiments were performed at least in triplicate and the data represented were the mean ± s.d. (**, *p*<0.01).

In an attempt to reveal the roles of miR-137 and miR-7847 in apoptosis regulation, mud crabs were injected with AMO-miR-137 and AMO-miR-7847. Flow cytometry analysis revealed that both AMO-miR-137 and AMO-miR-7847 induced higher apoptosis (in terms of percentage of apoptotic cells), compared with the controls (WT and AMO-NC) (Fig. 3E). Similarly, caspase 3/7 activity was significantly increased in AMO-miR-137 or AMO-miR-7847 injected groups, compared to the controls (Fig. 3F). To examine the participation of miR-137 and miR-7847 in exosome-mediated virus suppression, mud crabs were co-injected with either exosome-PBS, exosome-WSSV, exosome-WSSV and miR-137-mimic, or exosome-WSSV and miR-7847-mimic and WSSV. The results showed that WSSV copy number was significantly lower in the exosome-WSSV and WSSV co-injected group, compared with the other groups (*P<0.05*) (Fig. 3G). These data indicate that both miR-137 and miR-7847 might be controlled by the exosomes derived from WSSV infection, to promote replication of WSSV in hemocytes of mud crabs.

### Interactions between miR-137 and miR-7847 and their targeted genes

To reveal the pathways mediated by miR-137 and miR-7847, their target genes were predicted using the Targetscan and miRanda software. It was found that the apoptosis-inducing factor (AIF) was the target gene of both miR-137 and miR-7847 (Fig. 4A). To confirm this result, synthesized miR-137 and miR-7847 as well as EGFP-AIF-3’UTR-miR-137/-miR-7847 or mutant plasmids (EGFP-ΔAIF-3’UTR-miR-137/-miR-7847 were co-transfected into *Drosophila* S2 cells (Fig. 4B). The results showed that the fluorescence intensity in cells co-transfected with EGFP-AIF-3’UTR-miR-137 or EGFP-AIF-3’UTR-miR-7847 was significantly decreased compared with cells co-transfected EGFP-ΔAIF-3’UTR-miR-137 or EGFP-ΔAIF-3’UTR-miR-7847, respectively (Fig. 4C). This suggest that miR-137 or miR-7847 could interact with AIF.

**Fig 4.**
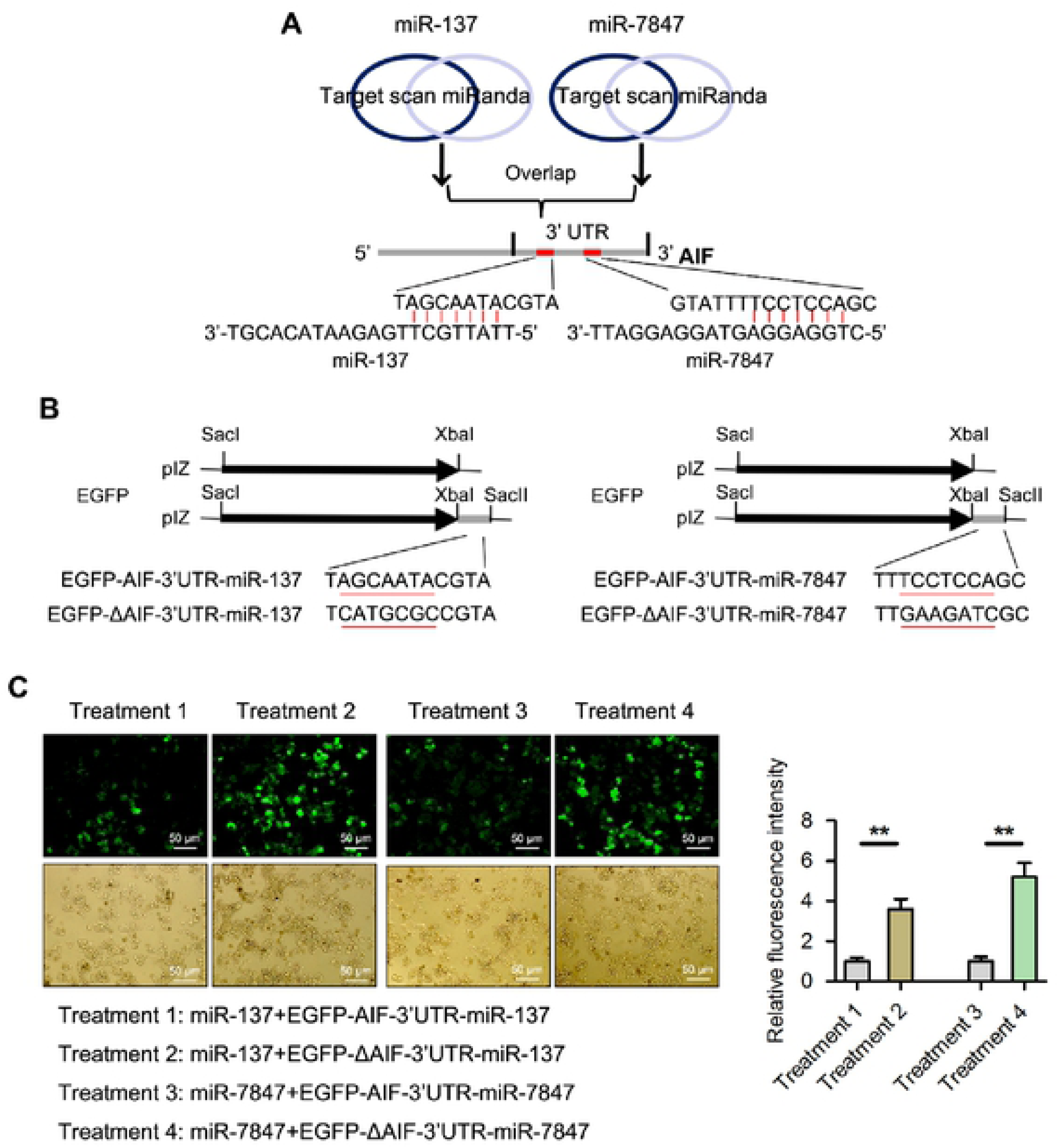
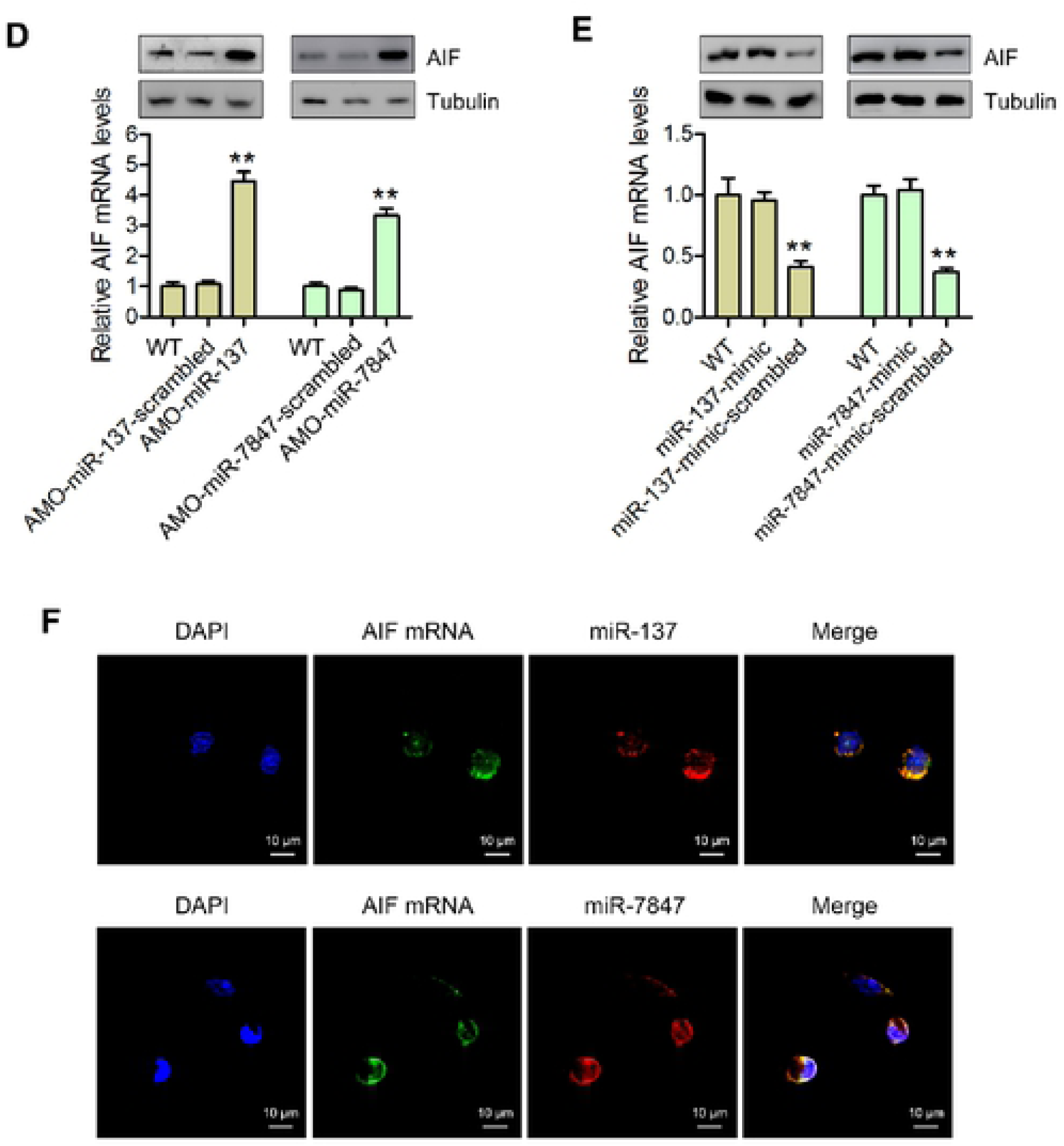
AIF is a direct downstream target for both miR-137 and miR-7847 in mud crab. **(A)** Target gene prediction of miR-137 and miR-7847 with two bioinformatics tools, as predicted, the 3’UTR of AIF could be simultaneously targeted by miR-137 and miR-7847. **(B)** The construction of the wild-type and mutated 3’UTRs of AIF. The sequences targeted by miR-137 and miR-7847 were underlined. **(C)** The direct interactions between miR-137, miR-7847 and AIF in insect cells, S2 cells were co-transfected with miR-137, miR-7847 and the indicated constructed plasmids for 48 h, then the relative fluorescence intensities were evaluated. **(D)** The effects of miR-137 and miR-7847 silencing on the expression levels of AIF in mud crab, AMO-miR-137 and AMO-miR-7847 were injected into mud crab separately, 48 h later, the mRNA and protein expression levels were examined. **(E)** The effects of miR-137 and miR-7847 overexpression on the mRNA and protein expression levels in mud crab. **(F)** The co-localization of miR-137, miR-7847 and AIF mRNA in mud crab hemocytes, miR-137, miR-7847, AIF mRNA and nucleus of hemocytes were respectively detected with FAM-labeled AIF mRNA probe (green), Cy3-labeled miR-137 and miR-7847 probe (red) and DAPI (blue). Each experiment was performed in triplicate and data are presented as mean ± s.d. (**, *p*<0.01)

In order to confirm the role of miRNAs in the expression of AIF mRNA, the expression of miR-137 and miR-7847 were silenced using AMOs or miRNA mimics and analyzed by qPCR. The qPCR results showed that AIF transcripts were significantly increased in following AMO-miR-137 or AMO-miR-7847 treatment, but was significantly decreased in the miR-137-mimic-scrambled or miR-7847-mimic-scrambled groups, respectively, compared with control (Fig. 4D and 4E). On the other hand, AMO-miR-137-scrambled or AMO-miR-7847-scrambled and miR-137-mimic or miR-7847-mimic had no significant effect on the expression of AIF mRNA (Fig. 4D and 4E).

To investigate the targeting of AIF by miR-137 and miR-7847 in hemocytes of mud crab, co-localization of miR-137/miR-7847 and AIF mRNA was examined. Hemocytes were treated with FAM-labeled AIF mRNA probe (green), Cy3-labeled miRNA probe (red) and DAPI (blue) before been observed under a fluorescence microscope. The results showed that miR-137/miR-7847 and AIF mRNA were co-localized in the cytoplasm of cells (Fig. 4F).

### Effects of AIF on WSSV proliferation and apoptosis

To ascertain whether AIF is involved in the immune response of mud crab, the expression profile of AIF was determined after WSSV challenge. The results (Western blot and qPCR analyses) revealed that AIF was significantly elevated at 24 and 48 hours post-WSSV challenge (Fig. 5A). In order to determine the effects of AIF on the proliferation of WSSV, viral copy number was examined in AIF depleted mud crabs challenged with WSSV. The results showed that WSSV copy number was significantly higher in AIF knockdown (siAIF-injected) mud crabs, compared with the control groups (GFP-siRNA and WSSV group) (Fig. 5B).

**Fig 5.**
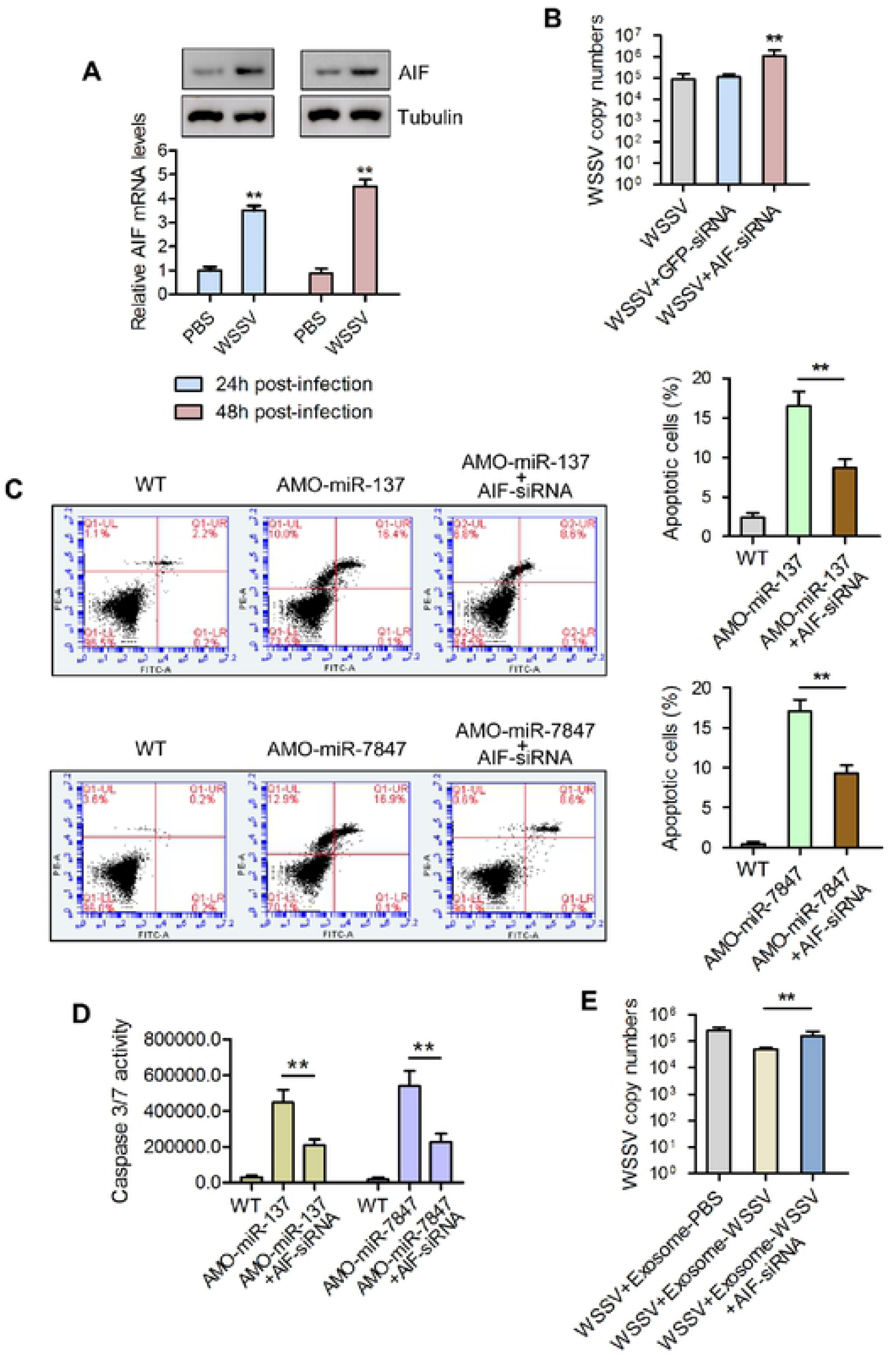
Exosomal miR-137 and miR-7847 regulate apoptosis and virus invasion through targeting AIF in mud crab. **(A)** The expression levels of AIF during WSSV infection, mud crabs treated with PBS or WSSV were subjected to western blot and qPCR analysis to detect the mRNA and protein levels of AIF. **(B)** The influence of AIF silencing on WSSV infection in mud crab. WSSV and AIF-siRNA were co-injected into mud crab for 48 h, followed by the detection of WSSV copy numbers, GFP-siRNA was used as control. (**C-D**) The involvement of AIF during miR-137 and miR-7847-mediated apoptosis regulation in mud crab. AMO-miR-137 and AMO-miR-7847 were co-injected with AIF-siRNA separately, then the hemocytes were subjected to annexin V assay **(C)** and caspase 3/7 activity analysis **(D)**. **(E)** The participation of AIF during exosome-mediated virus suppression, the indicated exosomes, WSSV and AIF-siRNA were co-injected into mud crabs, followed by the detection of WSSV copies. Data presented were representatives of three independent experiments (**, *p*<0.01).

In an attempt to unravel the roles of AIF in miR-137 and miR-7847-mediated apoptosis regulation, AIF-depleted mud crabs were injected with AMO-miR-137 or AMO-miR-7847. The results showed that the apoptotic rate and percentage of apoptotic cells were significantly higher in normal mud crabs injected with AMO-miR-137 or AMO-miR-7847 compared with AIF-depleted mud crabs injected with AMO-miR-137 or AMO-miR-7847, respectively (Fig. 5C). Similar results were obtained for caspase 3/7 activity analysis (Fig. 5D). Moreover, AIF was found to participate in exosome-mediated virus suppression. As shown in Fig. 5E, AIF-depleted mud crabs co-injected with exosome-WSSV and WSSV had significantly higher WSSV copy number compared with normal mud crabs co-injected with either exosome-WSSV or exosome-PBS and WSSV (*P<0.05*).

### Nuclear translocation of AIF induces DNA fragmentation

It has previously been reported that AIF is able to translocate into the nuclear of hemocytes [28]. Thus, we performed Western blot analysis to determine whether AIF was present in the nuclear extract of hemocytes from mud crabs injected with either WT (control), WSSV, AMO-miR-137 or AMO-miR-7847. The results revealed that AIF was found in the nucleus of hemocytes in all groups (except WT) at 6 and 24 hours post-injection (Fig. 6A). To further confirm the localization of AIF in the hemocytes of mud crabs, an immunofluorescent assay was carried out (Fig. 6B). The results indicated that AIF (detected using mouse anti-AIF antibody) was predominantly co-stained with DAPI in the nucleus of mud crab hemocytes. To explore the effect of AIF translocation to the nucleus, DNA fragmentation was analyzed using 3% agarose gel electrophoresis and genomic DNA isolated from hemocytes of mud crabs injected with WT (control), WSSV, AMO-miR-137 or AMO-miR-7847. The results revealed that WSSV, AMO-miR-137 and AMO-miR-7847 induced more DNA fragments compared with the control group (Fig. 6C).

**Fig 6.**
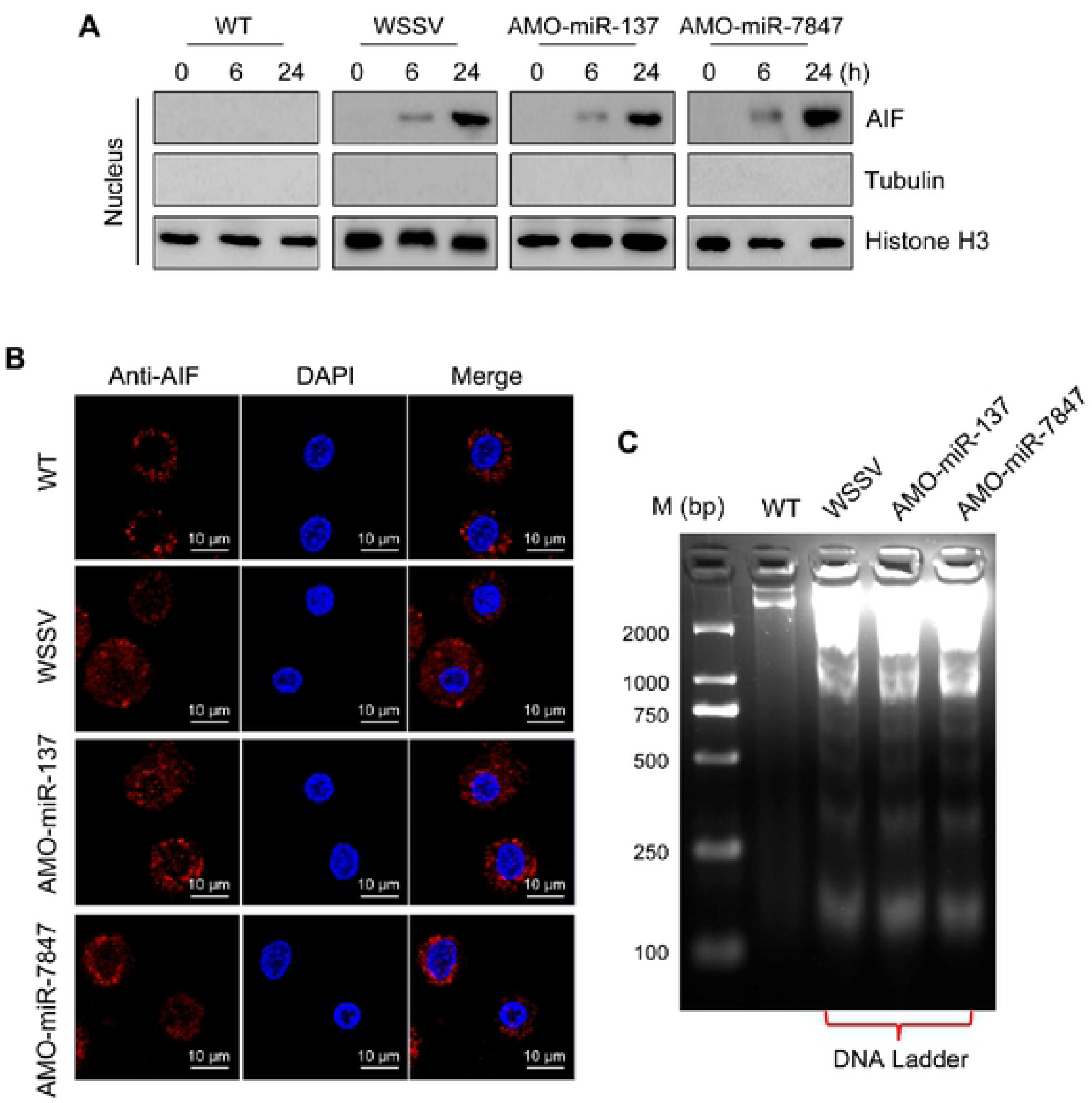
AIF translocated into nucleus to mediate DNA fragmentation. **(A)** The protein level of AIF in the nucleus, mud crabs were treated with WSSV, AMO-miR-137 and AMO-miR-7847 separately, at 0, 6 and 24 h after the treatments, the nucleus of hemocytes were isolated and subjected to western blot analysis. Tubulin and Histone H3 were used to evaluate the purity of the isolated nucleus. **(B)** Immunofluorescent assay was performed to detect the localization of AIF in mud crab hemocytes after specific treatment, including AMO-miR-137, AMO-miR-7847 and WSSV. Scale bar, 10 μm. **(C)** DNA Ladder detection of mud crab treated with AMO-miR-137, AMO-miR-7847 and WSSV separately, then the genomic DNA was isolated and subjected to agarose gel electrophoresis.

### Role of AIF in mitochondrial apoptosis

After observing that AIF was involved in hemocytes apoptosis, co-immunoprecipitation analysis was carried out, with the results (SDS-PAGE and Western blot) indicating that AIF could bind to HSP70 (Fig. 7A and 7B). To confirm the role of HSP70 in the regulation of apoptosis in hemocytes, mud crabs were depleted of HSP70 and the level of hemocytes apoptosis determined. It was observed that in hemocytes of HSP70 silenced (HSP70-siRNA) mud crabs, the apoptotic rate and percentage of apoptotic cells were significantly increased, compared with the control groups (WT and GFP-siRNA) (Fig. 7C). Knockdown of HSP70 also significantly decreased the WSSV copy number in hemocytes of mud crabs challenged with WSSV, compared with the control groups (Fig. 7D). This indicates the involvement of HSP70 in the proliferation of WSSV in mud crabs. Moreover, co-immunoprecipitation analysis revealed that HSP70 could bind to Bax (Fig. 7E and 7F). The interaction between HSP70 and Bax is shown in Fig. 7G. These results shows that the HSP70-Bax complex in hemocytes was disrupted when mud crabs were injected with either AMO-miR-137, AMO-miR-7847 or WSSV compared with control (WT group) (Fig. 7G).

**Fig 7.**
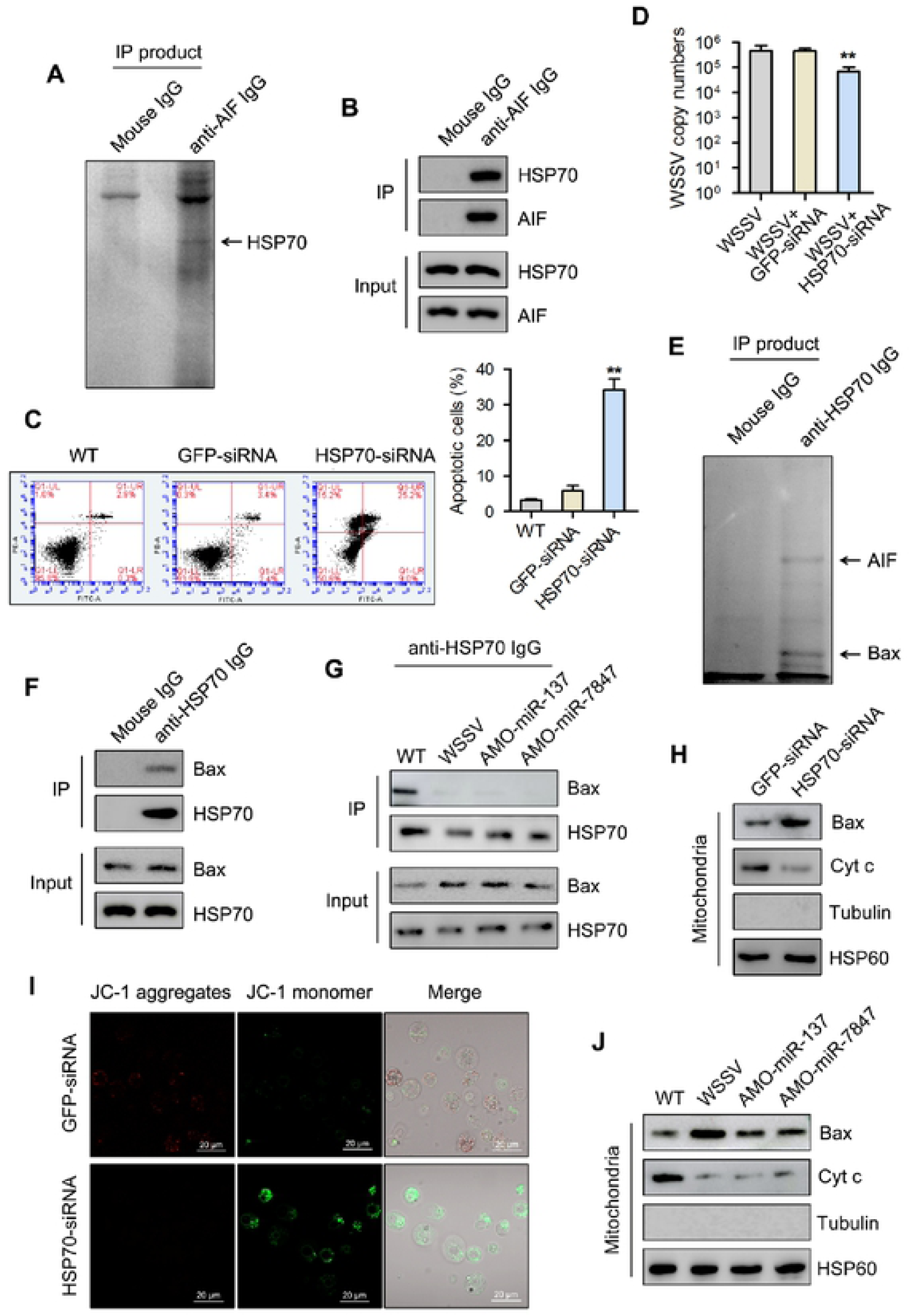
Underlying mechanisms of AIF-mediated mitochondrial apoptosis process. **(A)** Identification of proteins bound to AIF. The cell lysates of mud crab hemocytes were subjected to Co-IP assay using anti-AIF IgG, then the IP products were separated with SDS-PAGE and identified by mass spectrometry. **(B)** The interactions between AIF and HSP70 in mud crab, the cell lysates were subjected to Co-IP assay with anti-AIF IgG, then the IP products were subjected to Western blot analysis to detect HSP70. **(C)** The effects of HSP70 silencing on apoptosis regulation, HSP70-siRNA or GFP-siRNA were injected into mud crab for 48 h, then the hemocytes were subjected to annexin V assay. **(D)** The influence of HSP70 silencing on WSSV infection in mud crab. **(E)** Identification of proteins bound to HSP70, the proteins identified were indicated with arrows. **(F)** The interactions between HSP70 and Bax in mud crab, the cell lysates were subjected to Co-IP assay with anti-HSP70 IgG, then the IP products were subjected to Western blot analysis to detect Bax. **(G)** The interactions between HSP70 and Bax in mud crab with the indicated treatments. **(H)** The influence of HSP70 silencing on the protein levels of Bax and Cyt c in mitochondria. **(I)** The effects of HSP70 silencing on mitochondrial apoptosis, HSP70-siRNA or GFP-siRNA were injected into mud crab for 48 h, then the hemocytes were subjected to mitochondrial membrane potential measurement. **(J)** The detections of Bax and Cyt c in mitochondria of mud crab with the indicated treatments. All the numeral data represented the mean ± s.d. of triplicate assays (**, *p*<0.01).

In the mitochondria of HSP70-siRNA treated mud crabs, the expression of Bax was increased, but that of Cyt C decreased (Fig. 7H), which suggest a role of HSP70 in regulating the functions of Bax and Cyt C in the mitochondria. To further investigate whether silencing of HSP70 could affect mitochondrial mediated apoptosis, the mitochondrial membrane potential of hemocytes from HSP70-siRNA treated mud crabs was determined using confocal microscopy in terms of JC-1 aggregates (red fluorescence) and JC-1 monomers (green fluorescence) (Fig. 7I). The results (Fig. 7I) revealed weak red fluorescence and strong green fluorescence in the HSP70-siRNA treated group, compared with the GFP-siRNA control group, which indicates high apoptosis rate in HSP70 depleted mud crabs. To detect the presence of both Bax and Cyt C in mitochondria, mud crabs were injected with either WT (control), AMO-miR-137, AMO-miR-7847 or WSSV. The results (Western blot) revealed an increased level of Bax and a decreased level of Cyt C in mitochondria of the AMO-miR-137, AMO-miR-7847 or WSSV treated groups, compared with control (WT) (Fig. 7J).

Taken together, our findings revealed that during WSSV infection, both miR-137 and miR-7847 were less packaged in mud crab exosomes, then the decreased uptake of exosomal miR-137 and miR-7847 resulted in the activation of AIF in the recipient hemocytes. AIF could translocated to nucleus to induce DNA fragmentation, on the other hand, AIF competitively bind to HSP70 and led to the disintegration of HSP70-Bax complex, then the free Bax was transferred to mitochondria, which eventually caused mitochondrial apoptosis and suppressed virus infection in the recipient hemocytes (Fig. 8).

**Fig 8.**
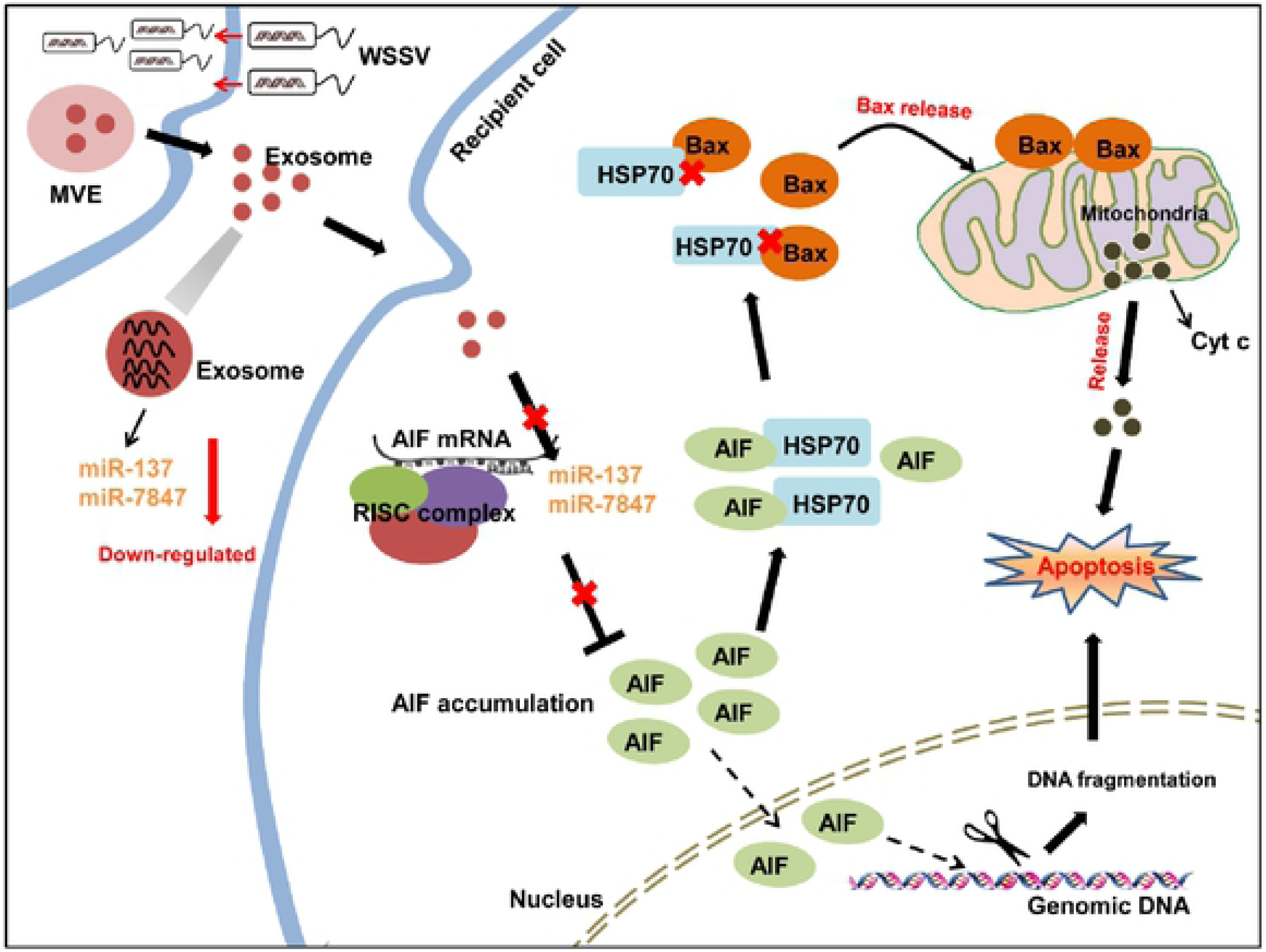
Proposed schematic diagram for the exosomal miR-137 and miR-7847-mediated apoptosis and virus invasion regulation in mud crab.

## Discussion

Exosomes are small membrane-enclosed vesicles actively released by cells into the extracellular environment, with the molecular cargo carried by exosomes reflecting the physiological or pathological state of donor cells [29]. In recent years, the involvement of exosomes in viral pathogenesis and immune responses has been widely investigated [8]. For instance, exosomes can protect viral contents from immune recognition, with studies showing that exosomes secreted by HCV-infected cells contain full-length viral RNA, which can be delivered to dendritic cells to establish infection [30, 31]. Meanwhile, VSV (vesicular stomatitis virus) infection can mediate the recruitment of TRAMP-like complex to exosomes to recognize and induce degradation of viral mRNAs [32]. In addition, exosomes released from HIV-infected cells contain regulatory factors required for apoptosis activation, which inhibit virus invasion by inducing apoptosis of uninfected cells [33]. Although there has been an increasing number of studies on exosomes and viral infection, most are focused on higher organisms, while the role of exosomes in antiviral immune response of invertebrates largely unexplored. In the current study, we revealed for the first time that, exosomes isolated from mud crab have typical characteristics as those in higher organisms. Furthermore, our data shows that exosomes released from WSSV-injected mud crabs could suppress virus invasion by inducing hemocytes apoptosis, which demonstrates a novel role of exosomes in invertebrates.

A distinct feature of exosomes is that a large number of nucleic acids are packaged in it, including miRNA, mRNA, mtDNA, piRNA, 1ncRNA, rRNA, snRNA, snoRNA and tRNA [34]. As the most abundant RNA in exosomes, studies have repeatedly demonstrated the essential roles of exosomal miRNAs during host-virus interactions [35]. Besides, the miRNA cargo carried by exosomes can be affected by external signals such as oxidative stress and pathogens infection [36], which possess completely different molecular composition to deal with these stimulations. For instance, EV71 infection cause differential packaging of miR-146a to exosomes, which suppresses type I interferon expression in the recipient cells, thus facilitating viral replication [37]. miR-483-3p is highly presented in mice exosomes during influenza virus infection, which potentiates the expression of type I interferon and proinflammatory cytokine genes [35]. Exosomal miR-145, miR-199a, miR-221 and Let-7f secreted by umbilical cord mesenchymal stem cells can directly bind to the genomic RNA of HCV and effectively inhibit viral replication [38]. In addition, studies have found that exosomal miRNAs are endowed with other functions apart from regulation of gene expression. Exosomal miR-21 and miR-29a secreted by HEK293 cells can serve as ligands and bind with toll-like receptors (TLRs), thus activating relevant immune pathways in the recipient cells [39]. Due to the diversity of their mode of regulation and functions, exosomal miRNAs are crucial regulators in response to virus infection, although no relevant research has been carried out in invertebrates. In the current study, we found that miR-137 and miR-7847 were less packaged in mud crab exosomes after WSSV challenge. Both miR-137 and miR-7847 are negative apoptosis regulators by targeting AIF, and thus decreased uptake of exosomal miR-137 and miR-7847 resulting in the activation of AIF, which eventually caused apoptosis and suppressed viral infection in the recipient hemocytes in mud crab. The present study thus reveals a novel molecular mechanism underlying the crosstalk between exosomal miRNAs and innate immunity in invertebrates.

The function of miR-7847 has not been reported previously, while miR-137 is regarded as an important regulator during tumorigenesis. It has been reported that miR-137 can inhibit the proliferation of lung cancer cells by targeting Cdc42 and Cdk6 [40]. Beside, miR-137 can also regulate the tumorigenicity of colon cancer stem cells through the inhibition of DCLK1 [41]. Thus far, the roles of miR-137 and miR-7847 in invertebrates had remained unclear, while our present study revealed that miR-137 and miR-7847 could suppress viral infection by promoting apoptosis. In addition, the present data revealed that AIF was co-targeted by both miR-137 and miR-7847 in mud crab. AIF is a mitochondrial FAD-dependent oxidoreductase protein that is involved in the regulation of oxidative phosphorylation [42]. Besides, AIF is the first identified caspase-independent protein important in mitochondrial pathway mediated apoptosis [43]. During the early process of the apoptosis, AIF is released from mitochondria and translocates to the nucleus [44], where it induces nuclear chromatin condensation, DNA fragmentation and cell death [45, 46]. It has been reported that when human alveolar epithelial cells (A549 cells) are challenged with influenza virus, AIF could translocate from mitochondria to nucleus, resulting in apoptosis in response to the virus infection [47]. At present, the function of AIF in the immunoregulation of invertebrate has not been addressed. Here, we found that AIF could inhibit WSSV infection by activating apoptosis of hemocytes in mud crabs. Moreover, AIF did not only translocate to the nucleus to induce DNA fragmentation, but was found to also competitively bind to HSP70 thereby disintegrating the HSP70-Bax complex, and freeing Bax, which transfers to the mitochondria to activate mitochondrial apoptosis pathway. The present study therefore provides some novel insight into the invertebrate innate immune system and highlights potential preventive and therapeutic strategies for viral diseases.

In summary, our findings revealed the evolutionary conservation of exosomal regulatory pathway in animals. As a topical research area, studies relevant to exosomes have largely focused on higher organisms [48]. In invertebrates, exosomes have been reported only in drosophila, and shown to be involved in the regulation of viral infection [49, 50] and miRNA biogenesis [51]. Thus, there is no enough evidence to conclude that exosome is a general regulatory approach in animals. In addition, exosomal miRNAs are also still unexplored in invertebrates, which means that there is an urgent need to characterize the existence and role of exosomal miRNAs. The current study was therefore the first to reveal the involvement of exosomal miRNAs in antiviral immune response of mud crabs, which shows a novel molecular mechanism of how invertebrates resist pathogenic microbial infection.

## Materials and Methods

### Mud crab culture and WSSV challenge

Healthy mud crabs, approximately 50 g each, were acclimated in the thanks filled with seawater at 25 °C for 3 days before WSSV challenge. To ensure that the crabs were virus-free before the experiments, PCR analysis were performed via WSSV-specific primer (5’-TATTGTCTCTCCTGACGTAC-3’ and 5’-CACATTCTTC ACGAGTCTAC-3’). Then, 200 μL WSSV (1×10^6^ *cfu*/ml) was injected into the base of the fourth leg of each crab. At different time post-infection, hemolymph was collected from three randomly chosen crabs per group for further investigation.

### Ethics statement

The mud crabs used in this study were taken from a local crab farm (Niutianyang, Shantou, Guangdong, China). No specific permits were required for the described field studies, as the sampling locations were not privately owned or protected in any way. Furthermore these field studies did not involve endangered or protected species. The animals were processed according to “the Regulations for the Administration of Affairs Concerning Experimental Animals” established by the Guangdong Provincial Department of Science and Technology on the Use and Care of Animals.

### Isolation and analysis of exosomes

For exosomes isolation, the hemolymph of mud crabs was separated, after centrifuged at 300 × g for 5 min, the sediment was removed. Subsequently, the supernatant was subjected to ultracentrifugation, followed by sucrose density-gradient centrifugation and filtrated through 0.22 μm filters. Then the obtained exosomes were observed by Philips CM120 BioTwin transmission electron microscope (FEI Company, USA) and quantified by Nano-Sight NS300 (Malvern Instruments Ltd, UK).

### Exosomes tracing

For exosome-tracing experiments, the isolated exosomes were pre-treated by DiO (Beyotime, China) and injected into mud crabs for 2 h. Then, the hemocytes were isolated and treated with DiI (Beyotime), followed by observation with confocal laser scanning microscopy TCS SP8 (Leica, Germany).

### Microarray analysis of exosomal miRNAs

Exosomal miRNAs microarray analysis was performed at Biomarker Technologies (Beijing, China), using Agilent Human miRNA 8*60 K V21.0 microarray (Agilent Technologies, USA). The Gene Expression Omnibus accession number is PRJNA600674. Gene Spring Software 12.6 (Agilent Technologies) was used for quantile normalization and data processing. Besides, Pearson’s correlation analysis through Cluster 3.0 and TreeView software was used for Hierarchical clustering analysis of the differential expression of miRNAs.

### RNA interference assay

Based on the sequence of *Sp*-AIF (GenBank accession number MH393923.1) and *Sp*-HSP70 (GenBank accession number EU754021.1), the siRNA specifically targeted *Sp*-AIF or *Sp*-HSP70 gene was designed, generating AIF-siRNA (5’- UCUAAUUCUGCAUUGACUCUGUU-3’) and HSP70-siRNA (5’- UCUUCAUAA GCACCAUAGAGGAGUU-3’). The siRNAs were synthesized using *in vitro* Transcription T7 Kit (TaKaRa, Dalian, China) according to the user’s instructions. Then, 50 μg AIF-siRNA or HSP70-siRNA was injected into each mud crab respectively. At different time post siRNA injection, three mud crabs were randomly selected for each treatment and stored for further use.

### Quantification of mRNA with real-time PCR

The real-time quantitative PCR was conducted with the Premix Ex Taq (Takara, Japan) to quantify the mRNA level. Total RNA was extracted from hemocytes, followed by first-strand cDNA synthesis using PrimeScript™ RT Reagent Kit (Takara, Japan). Primers AIF-F (5’-AGCCATTGCCAGTCTTTGAT-3’) and AIF-R (5’-GAACCCAGAAATCCTCCACC-3’) was used to quantify the AIF mRNA transcript, while primers β-actin (β-actin-F, 5’-GCGGCAGTGGTCATCTCCT-3’ and β-actin-R, 5’-GCCCTTCCTCACGCTATCCT-3’) was used to quantify the internal control β-actin. Relative fold change of mRNA expression level of AIF was determined using the 2^-ΔΔCt^ algorithm [52].

### Quantification of miRNA with real-time PCR

Total RNA was extracted using MagMAX^TM^ mirVana^TM^ Total RNA Isolation Kit (Thermo Fisher Scientific, USA), followed by first-strand cDNA synthesis via PrimeScript^TM^ II 1st Strand cDNA Synthesis Kit (Takara, Japan) using miR-137- primer(5’-GTCGTATCCAGTGCAGGGTCCGAGGTCACTGGATACGACACGTG TAT-3’) and miR-7847-primer (5’- GTCGTATCCAGTGCAGGGTCCGAGGTCACTG GATACGACAATCCTCC-3’). Real-time PCR was carried out with the Premix Ex Taq (Takara, Japan) to quantify the expression level of miR-137 and miR-7847, U6 was used as control, the primers used were listed below. miR-137-F (5’- CGCCGTTATTGCTTGAGA-3’) and miR-137-R (5’- TGCAGGGTCCGAGGTCAC TG-3’), miR-7847-F (5’-CGCCGCTGGAGGAGTAGG-3’) and miR-7847-R (5’- TGCAGGGTCCGAGGTCACTG-3’), U6-F (5’-CTCGCTTCGGCAGCACA-3’) and U6-R (5’-AACGCTTCACGAATTTGCGT-3’).

### Analysis of WSSV copies by real-time PCR

The genomic DNA of WSSV-infected mud crab was extracted using a SQ tissue DNA (Omega Bio-tek, Norcross, GA, USA) according to the manufacturer’s instruction. To detect WSSV copies in mud crab, real-time PCR analysis was carried out using Premix Ex Taq (probe qPCR) (Takara, Dalian, China). Real-time PCR was performed with WSSV-specific primers WSSV-RT1 (5’-TTGGTTTCATGCCCGAGA TT-3’) and WSSV-RT2 (5’-CCTTGGTCAGCCCCTTGA-3’) and a TaqMan probe (5’-FAM-TGCTGCCGTCTCCAA-TAMRA-3’) according to previous study [53]. The internal standard of real-time PCR was a DNA fragment of 1400 bp from the WSSV genome [54].

### Detection of apoptotic activity

In order to evaluate the apoptotic activity of mud crab, the Caspase 3/7 activity of hemocytes was determined with the Caspase-Glo 3/7 assay (Promega, USA). While the apoptosis rate was evaluated using FITC Annexin V Apoptosis Detection Kit I (BD Pharmingen^TM^, USA) according to the manufacturer’s instruction. And flow cytometry (AccuriTM C6 Plus, BD biosciences, USA) was used to analyze the apoptosis rate. Besides, the mitochondrial membrane potential, an indicator of the apoptotic activity in hemocytes, which were measured by Mitochondrial membrane potential assay kit with JC-1 (Beyotime, China) following the protocols and finally analyzed by confocal microscope(ZEISS, Germany).

### Western blot analysis

The hemocytes of mud crab were homogenized with RIPA buffer (Beyotime Biotechnology, China) containing 1 mM phenylmethanesulfonyl fluoride (PMSF) and then centrifuged at 13,000×g for 5 min at 4 °C. Then the cell extracts mixed with 5 × SDS sample buffer were separated by 12 % SDS-polyacrylamide gel electrophoresis and transferred onto a nitrocellulose membrane (Millipore, USA). Subsequently, the membrane was incubated in blocking buffer (Tris-buffered saline containing 0.1% Tween 20 (TBST) and 5% (W/V) nonfat dry milk) and further incubated with appropriate primary antibody at 4 °C. After washed with TBST, the membrane was incubated with horseradish peroxidase-conjugated secondary antibody (Bio-Rad, USA) for subsequent detection by ECL substrate (Thermo Scientific, USA).

### The target gene prediction of miRNA

Targetscan (http://www.targetscan.org) and miRanda (http://www.microrna.org/) were used to predict the target genes of miR-137 and miR-7847 by a commercial company (BioMarker, Beijing, China). And the overlapped target gene predicted by the two algorithms were the candidate target gene.

### Cell culture, transfection, and fluorescence assays

The *Drosophlia* Schneider 2 (S2) cells were cultured in Express Five serum-free medium (SFM) (Invitrogen, USA) at 27 °C. The EGFP-AIF-3’UTR or the mutant plasmids (100 ng/well) and the synthesized miR-137 or miR-7847 (50 nM/well) were co-transfected into S2 cells using with Cellfectin II Reagent (Invitrogen, USA) according to the manufacturer’s protocol. At 48 h after co-transfection, the EGFP fluorescence in S2 cells was measured by a Flex Station II microplate reader (Molecular Devices, USA) at 490/ 510 nm of excitation/emission (Ex/Em).

### The silencing or overexpression of miR-137 and miR-7847 in mud crab

Anti-microRNA oligonucleotide (AMOs) or miRNA mimic was injected at 30 μg/crab to knockdown or overexpress miRNAs in mud crab, AMO-miR-137 (5’- ACGTGTATTCTCAAGC**A**AT**A**A-3’), AMO-miR-7847 (5’-AATCCTCCTACTCC**T** CC**A**G-3’), miR-137 mimic (5’-TTATTGCTTGAGAATA**C**AC**G**T-3’) and miR-7847 mimic (5’-CTGGAGGAGTAGGA**G**GA**T**T-3’) were modified with 2’-O-methyl (OME) (bold letters) and phosphorothioate (the remaining nucleotides). All oligonucleotides were synthesized by Sangon Biotech (Shanghai, China). At different time points after the last injection, three mud crabs per treatment were collected for subsequent use.

### Fluorescence *in situ* hybridization

The hemocytes of mud crab were seeded onto the polysine-coated cover slips, fixed with 4% polyformaldehyde for 15 min at room temperature. After that, the cover slips were dehydrated in 70% ethanol overnight at 4°C, followed by incubation with hybridization buffer [1× SSC (15 mM sodium citrate, 150 mM sodium chloride, pH 7.5), 10% (w/v) dextran sulfate, 25% (w/v) formamide, 1× Denhardt’s solution] containing 100 nM probe for 5 h at 37°C. The miR-137 probe (5’-FAM- ACGTGTATTCTCAAGCAATAA-3’), miR-7847 probe (5’-FAM- AATCCTCCTACT CCTCCAG-3’) and AIF probe (5’-Cy3-TCCATCTTCTGTACTCTTGACT-3’) were used. Then the slips were washed with PBS for three times, and after that the hemocytes were stained with DAPI (4’, 6-diamidino-2-phenylindole) (50 ng/ml) (Sigma, USA) for 5 min [55]. The images were captured using a CarlZeiss LSM710 system (Carl Zeiss, Germany).

### Statistical analysis

All data were subjected to one-way ANOVA analysis using Origin Pro8.0, with *P* <0.01 considered statistically significant. All assays were biologically repeated for three times.

## Acknowledgements

This study was financially supported by the National Natural Science Foundation of China (31802341, 41876152), Natural Science Foundation of Guangdong Province, China (2018A030307044), Department of Education of Guangdong Province, China (2017KQNCX072), STU Scientific Research Foundation for Talents (NTF18001), and Guangdong Provincial Special Fund for Modern Agriculture Industry Technology Innovation Teams (2019KJ141).

## Author contributions

YG and TTK performed the experiments and analyzed the data. SKL and YG wrote the manuscript. All authors read and approved the contents of the manuscript and its publication.

